# Canine major histocompatibility complex class I (MHC-I) diversity landscape

**DOI:** 10.1101/2024.02.14.580220

**Authors:** Yuan Feng, Kun-Lin Ho, Mengyuan Zhang, Nikitha Brahmasamudra Sundaresha, Hannah Lorelle Cavanagh, Shaying Zhao

## Abstract

The genes of the Major Histocompatibility Complex class I (MHC-I) are among the most diverse in the mammalian genome, playing a crucial role in immunology. Understanding the diversity landscape of MHC-I is therefore of paramount importance. The dog is a key translational model in various biomedical fields. However, our understanding of the canine MHC-I diversity landscape lags significantly behind that of humans. To address this deficiency, we used our newly developed software, KPR *de novo* assembler and genotyper, to genotype 1,325 samples from 1,025 dogs with paired-end RNA-seq data from 43 BioProjects, after extensive quality control. Among 926 dogs that pass the QC, 591 dogs (64%) have at least one allele genotyped, and a total of 97 known alleles and 52 putative new alleles were identified. Further analysis reveals that DLA-I gene expression levels vary among the tissues, with lowest for testis and brain tissues and highest for blood, corpus luteum, and spleen. We identified dominant alleles in each of the 17 canine breeds, as well as among the entire canine population. Furthermore, our analysis also identifies breed-specific alleles and mutually co-occurred/exclusive alleles. Our study indicates that canine DLA-88 is as diversified as human HLA-A/B/C genes within the entire population, but less diversified within a breed than with HLA-A/B/C within an ethnic group. Lastly, we examined the hypervariable regions (HVR) within or across human/canine MHC-I alleles and found that 80% of the HVRs overlap between the two species. We further noted that 80% of the HVRs are within 4A contact with the peptides, and that the dog-human difference overlaps with only 20% HVRs. Our research offers valuable insights for immunological studies involving dogs.

## Introduction

As a mammalian species with intact immune system ^1^, spontaneous cancer, and has high sequence homology to human genome, dog has been used as an animal model for drug discovery, oncology research, and immunotherapy in human for years. Moreover, most of the dogs share the same environments as humans after domestication and are exposed to similar carcinogens and pathogens. However, due to a lack of resources, canine oncology and immunology research is far behind human, including the understanding of its major histocompatibility complex class I (MHC-I) molecules.

The MHC-I molecules are encoded by the most polymorphic regions in mammalian genome. And functionally, these molecules are expressed on the surface of all nucleated cells and play an essential role in presenting and exposing intracellular antigens to immunological surveillance ^2^. These antigens may come from viral infections, or self-peptides undergone somatic alterations in tumorigenesis ^3^. Therefore, understanding the MHC-I diversity at individual level is crucial. In human, some specific MHC-I alleles show higher efficiency against HIV ^4^. And the T cell receptor (TCR) repertoire is also associated with MHC-I diversity ^5^, which reflects the features of the immune system.

Meanwhile, it is also important to study MHC-I from the population aspect. In human, a total of 23,417 alleles have been identified so far from the three classical MHC-I encoding genes ^6^. However, some alleles show significantly higher frequencies than the rest, especially in specific ethnic groups. In Greece population, the dominant HLA-A alleles are A*02 and A*24, and B*51 and B*18 for HLA-B gene ^7^. But in a Chinese population studied in Guizhou, the most frequent alleles are A*11 and A*02 for HLA-A, and B*40 and B*46 for HLA-B ^8^. These frequencies are important for MHC-I related disease research. Some dominant alleles should be prioritized in the studies. And confirming the association between MHC-I genotypes and diseases is critical in predicting the therapeutic outcomes in different groups. Canine faces the same issue. Some breeds are more susceptible to some specific diseases, for instance, the high occurrences of lymphoma, hemangiosarcoma and osteosarcoma in Golden Retriever ^9^. And it has been reported that canine hypoadrenocorticism is associated with MHC class II molecules ^10^. Our previous studies identified prevalent MHC-I alleles in Maltese, Poodle, Yorkshire Terrier, and Shih Tzu ^11^. And identifying high-frequency alleles will be the first step to dig into the relationship between MHC-I and these diseases.

Large scale MHC-I diversity landscape study was hampered previously by the lack of a high-throughput method, and the absence of an *in silico* approach to identify new canine MHC-I alleles. This problem was solved using the Kmer-based paired-end read (KPR) *de novo* assembler and genotyper ^11^. Meanwhile, the pair-end RNA-seq data from over 1,400 samples is publicly available in Sequence Read Archive (SRA).

The aims of this study include the identification of new allele candidates, and prevalent alleles in major dog breeds and population. We also analyzed the transcription rates across different tissue types, age groups and genders, as it is a determinant of MHC-I synthesis, and directly influencing the crosstalk efficiency to the CD8^+^ T cells and the therapeutic outcomes. Using the newly identified alleles together with the known ones, we refined current canine MHC-I allele classification based on hyper-variable regions (HVR) ^12^.

## Results

### Quality Control (QC) of Published Canine Sequencing Data

The RNA-seq dataset comprises 1,325 samples obtained from 1,025 animals across 43 BioProjects (Table S1). These encompass diverse cancer types, including 161 mammary tumors (MT)^13,14^, 66 B-cell lymphomas (BCL), 15 T-cell lymphomas (TCL), 89 osteosarcomas (OSA)³, 76 oral melanomas (OM), 74 hemangiosarcomas (HSA)^15,16^, 39 gliomas (GLI)_, 26 bladder carcinomas (TCC), 11 prostate carcinomas (PRO)^17,18^, 10 oral squamous cell carcinomas (OSCC), 2 melanomas (MEL), along with other non-tumor samples (refer to Table S1). The RNA-seq data were generated by different groups. We hence performed a rigorous QC to ensure that data chosen from each study meets a set of quality standards before any integrative analysis.

Regarding sequencing amount, all datasets exhibit a median ranging from 15 million (M) to 90 M read pairs per sample (Fig. 1a; Table S1). With all but one sample having >5_M read pairs, we proceeded to evaluate the mapping of read pairs to the canine reference genome³. Except for three BioProjects, nearly all studies demonstrated >30% read pairs uniquely and concordantly mapped to the genome, with the median close to or greater than 40% (Fig. 1b). Samples with uniquely and concordantly mapping rates <30% (35 samples) were excluded (Table S1). In addition, 11 samples with low mapping quality were excluded (Fig. 1c; Table S1). For the CDS target rate, the majority of samples exhibited, on average, >50% read pairs that were uniquely and concordantly mapped to the CDS regions (Fig. 1d; Table S1). 15 samples with poor CDS target rates were excluded. The overall distribution of expression levels of ∼200,000 protein-coding genes in each sample was also examined, identifying three significant outliers (Fig. 1e; Table S1). Following these rigorous quality control processes, 1,258 BioSamples and 926 dog samples successfully passed for further analysis.

**Figure 1.**
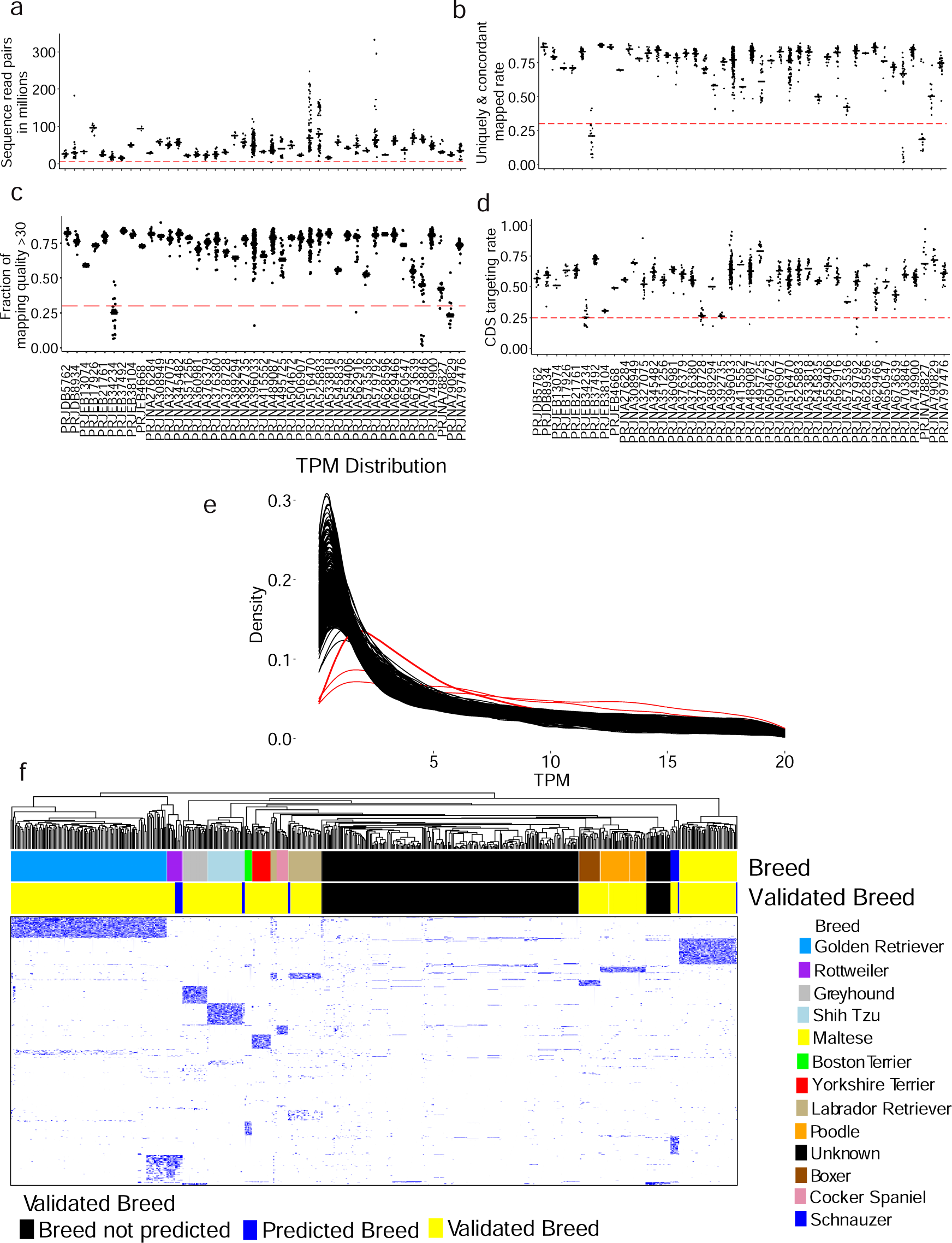
We performed a rigorous quality control (QC) on 1325 canine RNA-seq samples from 43 BioProjects. a. Distributions of total read pairs per sample across each study, with each dot representing a sample and the median highlighted by a black line. The dashed line denotes the QC cutoff. Each study is represented by the BioProject. b-d. Distributions of per-sample rate of read pairs that aligned concordantly and uniquely to the canFam3 reference genome (b), fractions of reads with mapping quality of ≥30 (c), CDS targeting rate (the fraction of read pairs that align concordantly and uniquely to the canFam3 CDS regions) (d). e. Distribution of TPM values of 1325 samples. Three red lines indicate the outliers and were removed from further analysis. f. Breed validation and prediction using breed-specific germline base substitutions and small indels discovered as described ^9^. The heatmap visualizes the clustering of 587 animals, comprising 347 dogs with provided breeds and 240 dogs without provided breeds. The clustering is based on variant allele frequency (VAF) values derived from the analysis of 2,813 breed-specific germline base substitutions and small indel variants. The “Provided Breed” bar indicates the breed of each dog provided by the source studies. The “Validated Breed” bar denotes the analysis outcome as specified. See also Table S1.

To validate breed data accuracy, focus was directed toward the 12 pure breeds in the RNA-seq dataset, each meeting QC measures specified in Fig. 1(a-e). Breed-specific germline variants were identified as described in^9^, yielding 2,183 breed-specific variants (Table S1). These variants, characterized as germline base substitutions and small indels unique to or enriched in specific breeds, were subjected to clustering analysis based on variant allele frequency (VAF) values. The analysis confirmed the breeds of 347 dogs and predicted the breed for 12 dogs without a breed label (Fig. 1f). These dogs were subsequently employed for downstream breed-related analyses, as described later.

### New DLA-I Alleles Identification

In our previous study, we introduced a software named KPR^11^ designed for the identification of MHC-I alleles in canine (DLA-I) paired-end RNA-seq data. Leveraging this software, we applied it to publicly available datasets. KPR successfully identified 2,315 contigs from 612 dogs in total 1025 dogs, achieving a 60% genotyping rate. To validate our allele identification, we incorporated five key features, as presented in the Venn diagram (Fig. 2a): Relative expression >25% (REL >25%), ranking among the top four highest relative expression levels within a sample (Top4), presence in ≥3 dogs (Freq ≥ 3), designation as known alleles (Known), or validation through Seq2HLA^19,20^. Notably, the majority of contigs exhibited features from more than two categories, with 461 contigs possessing all specified properties (Fig. 2a). For new allele identifications, contigs with at least one property or those reported in^21^ were considered validated alleles. Among all validated new alleles, we identified 32 from DLA-88, 4 from DLA-88L, 10 from DLA-12, 4 from DLA-64, and 2 from DLA-79. A comprehensive summary of DLA-I allele identifications is presented in Fig. 2b. To further elucidate the relationships among identified alleles, a phylogenetic tree encompassing 149 different alleles was reconstructed using amino acid sequences from exon2 and exon3. The resulting tree distinctly categorized DLA-88, DLA-12, DLA-64, and DLA-79 alleles into four lineages, with DLA-64 and DLA-79 forming outlier lineages. Noteworthy is the cohesive clustering of alleles within their respective gene lineages, except for three DLA-12 alleles, which clustered with the DLA-88 lineage due to high amino acid sequence similarity. All DLA-88L alleles were inclusively placed within the DLA-88 lineage (Fig. 2c).

**Figure 2.**
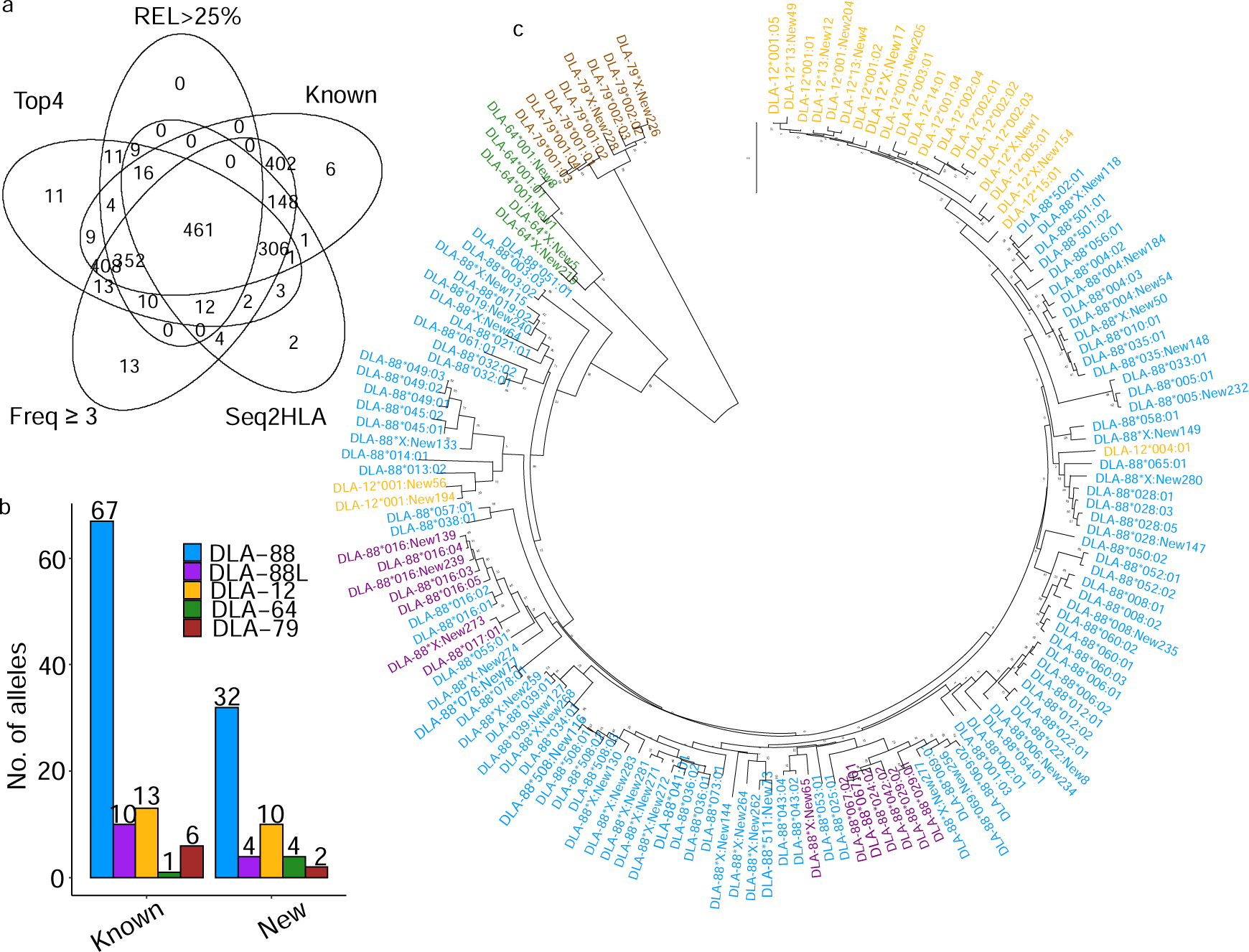
We identified a total of 52 new DLA-I alleles from 612 dogs. a. Number of contigs identified in 612 Dogs: Each number represents the count of contigs meeting criteria such as relative expression > 25% (REL > 25%), contigs in one of the top 4 highest relative expression levels within a sample (Top4), presence in ≥ 3 dogs (Freq ≥ 3), known alleles (Known), or validated with Seq2HLA (Seq2HLA). A total of 461 contigs exhibit all these properties. b. Distributions of the known and new allele numbers from different DLA genes in 612 dogs. Different allele genes are represented in different colors. DLA-88: blue, DLA-12: yellow, DLA-88L: purple, DLA-64: green, DLA-79: brown. c. Neighbor-joining phylogenetic tree with exon 2–exon 3 amino acids sequences of 145 alleles of DLA-88, DLA-12, DLA-88L, DLA-64 and DLA-79 alleles. The color for each allele gene is represented as b. Newly identified alleles in this study are presented as bold.

### Factors Affecting Genotyping Rates

In the present study, our genotyping software, KPR, did not achieve a complete genotyping rate of 100%. Subsequently, an analysis was conducted to discern the underlying factors influencing genotyping rates. The findings revealed a pronounced association between successfully genotyped samples and elevated read pairs within exon2 and exon3 regions, coupled with diminished sequence error rates compared to samples that failed genotyping (Fig. 3a and 3b). Interestingly, the investigation into sequence evenness unveiled no significant impact on genotyping rates (refer to Fig. 3c). This observation underscores the importance of high sequence coverage and low sequence error rates in achieving reliable genotyping outcomes.

**Figure 3.**
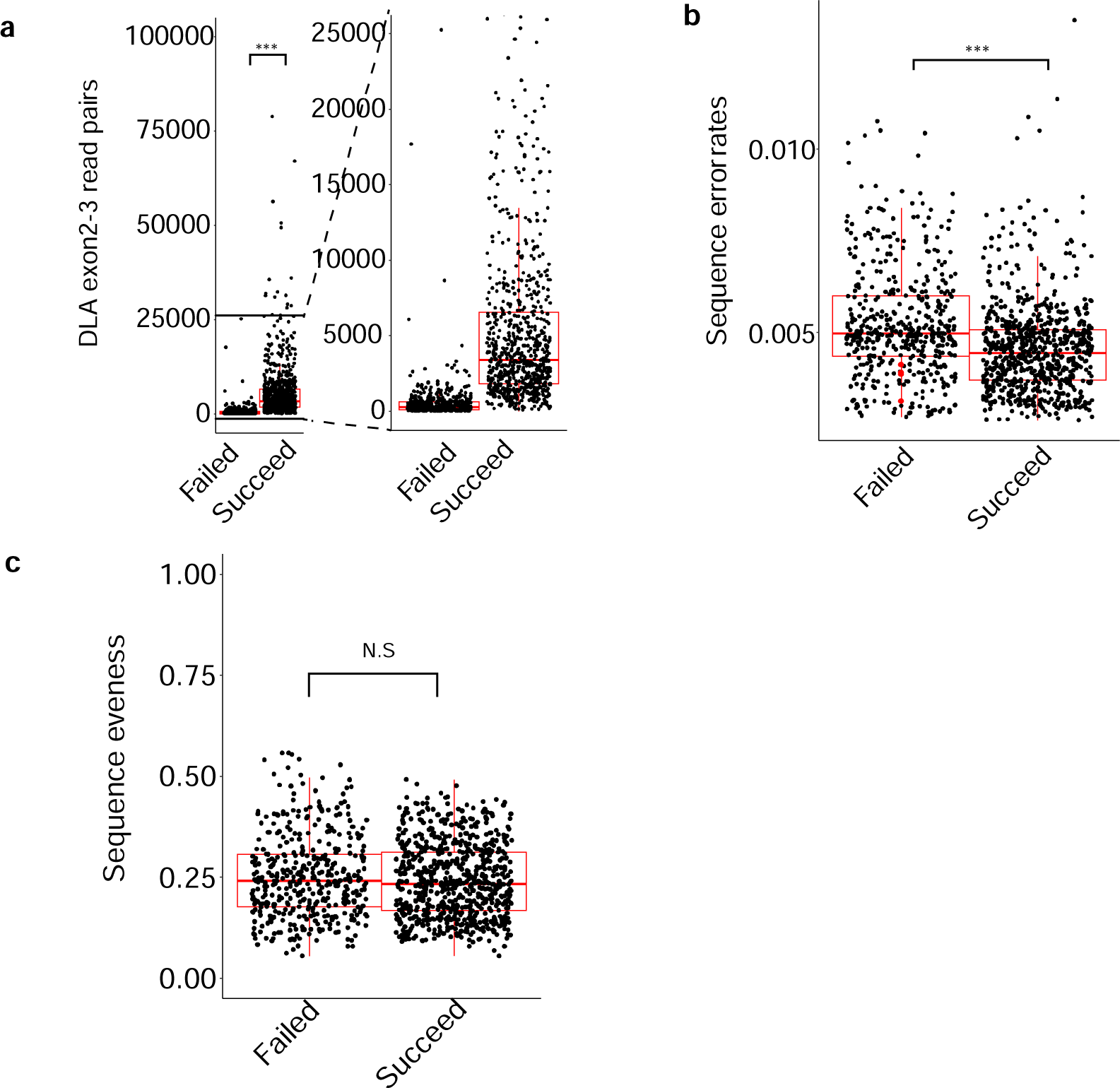
Genotyping is influenced by DLA read pairs and sequence error rates. a. Distribution of DLA exon2-3 reads in successfully and unsuccessfully genotyped samples: Each dot represents a sample, with the median indicated by a red line. Box plots depict the median, 25th, and 75th percentiles (*** indicating p = 2e−16). b. Distribution of sequence error rates in successfully and unsuccessfully genotyped samples: Box plots display the median, 25th, and 75th percentiles (*** indicating p = 2e−16). c. Distribution of sequence evenness rates in successfully and unsuccessfully genotyped samples: Box plots display the median, 25th, and 75th percentiles (NS indicating p = not significant).

### Tissue-Specific Expression Patterns of the DLA-I Genes

Our research aimed to elucidate tissue-specific and tumor-type dependent expression patterns of the DLA-I gene. This effort sought to uncover gene expression across diverse tissues and tumor types. Across all tissues examined, a consistent pattern emerged, revealing high expression levels of all DLA genes in the spleen, and lower expression levels in the testis and brain (Fig. 4, left panel). Notably, while different DLA genes exhibited similar trends across different tissues, DLA-88 and DLA-12 consistently exhibited high expression levels, but DLA-64 and DLA-79 consistently exhibited low expression levels (Fig. 4b, left panel). Noteworthy, DLA-79 exhibited elevated expression levels in muscle tissue, despite its relatively lower expression levels across other tissues. Conversely, although DLA-88 and DLA-12 exhibited high overall expression levels, their expression in muscle tissue remained comparably low (Fig. 4a, left panel). Furthermore, our investigation extended to exploring the differential expression of DLA-I genes across different tumor types. We observed the highest expression levels in T-cell and B-cell lymphomas (TCL, BCL), while gliomas (GLI) and bladder carcinomas (TCC) displayed comparatively lower expression levels (Fig. 4, right panel). These findings underscore the intricate interplay of tissue-specific and tumor-type dependent gene expression patterns within the DLA-I gene family, emphasizing the need for further research to elucidate their functional implications.

**Figure 4.**
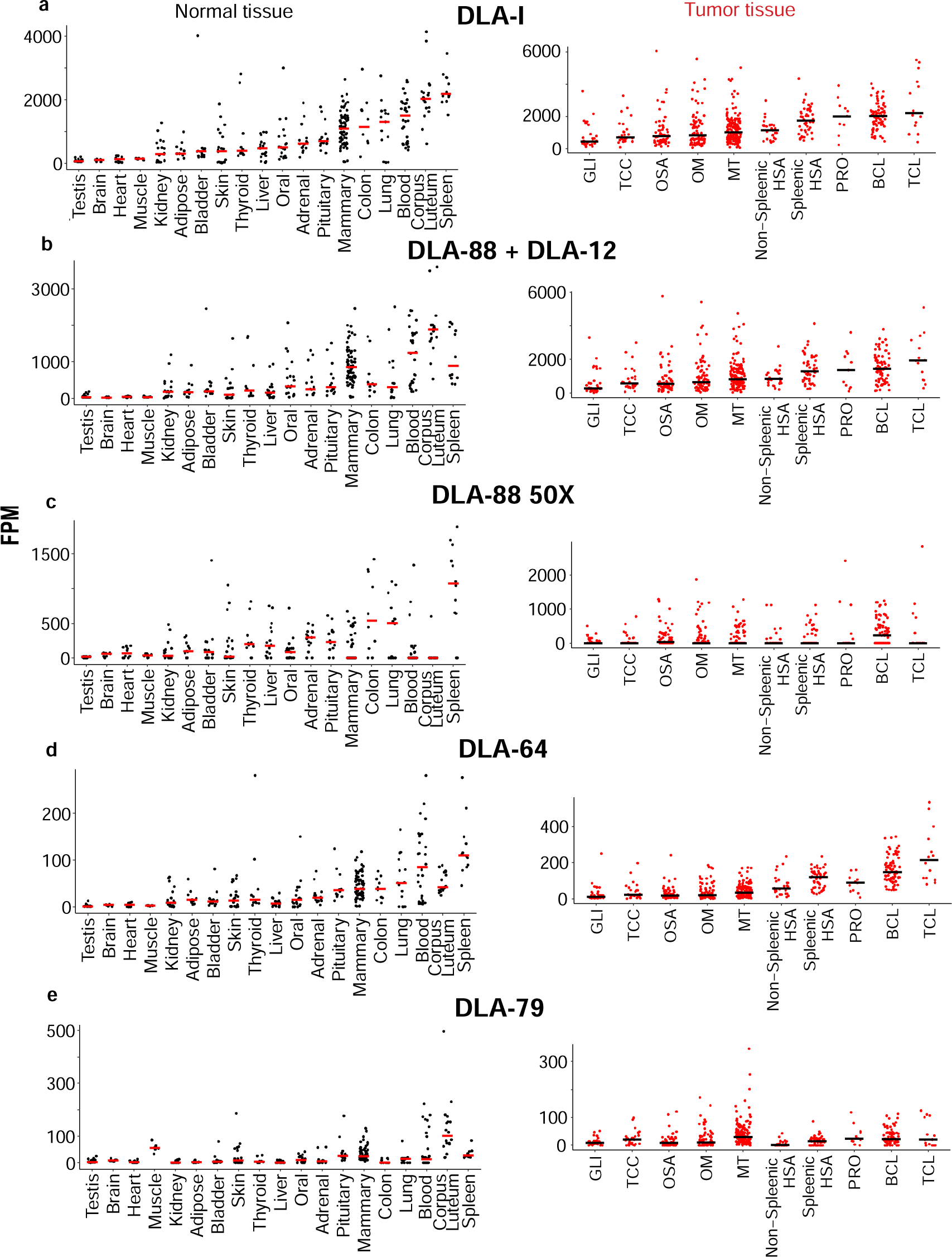
DLA-I expression across different tissues and tumor types. a. FPM distributions of DLA-I per sample for each tissue, arranged from left to right based on ascending median values, with subsequent plots adhering to the same order. Each black dot represents a sample of normal tissue, and the red line indicates the median value. Each red dot represents a tumor sample, and the black line indicates the median value. In the tumor samples, each study is represented by the tumor type. MT mammary tumor, GLI glioma, BCL B-cell lymphoma, TCL T-cell lymphoma, OM oral melanoma, OSA osteosarcoma, HSA hemangiosarcoma, PRO prostate carcinoma, TCC bladder carcinoma. Tissues with more than 8 samples after our QC are shown in the figures. See also FigureS4. b-e. FPM distributions of DLA-88+DLA-12 (b), DLA-88 50X (c), DLA-64 (d), and DLA-79 (e) per sample of each tissue and tumor type.

### Different Breeds Demonstrate Varying Dominant DLA-I Alleles

Research has established that different ethnicities can exhibit distinct patterns of dominant MHC-I allele expression. We sought to explore whether a similar relationship exists between dog breeds and MHC-I allele expressions. Among all genotyped dogs, we observed that DLA-88*501:01 and DLA-88*012:01 are prominently expressed within the DLA-88 locus, with several dominant alleles evident. Similarly, highly expressed DLA-88L alleles exhibit comparable expression levels to DLA-88 and display multiple dominant alleles. In contrast, DLA-12, DLA-64, and DLA-79 tend to feature one highly expressed allele, indicating higher diversity within DLA-88 and DLA-88L than in other DLA genes (Fig. 5a). Furthermore, our investigation revealed that different breeds tend to manifest distinct dominant alleles. Specifically, DLA-88*051:01 and DLA-88*038:01 emerge as the top two highest expression alleles in Golden Retrievers (Fig. 5b). Despite the prevalence of golden retrievers in the dataset, we did not observe these same top two dominant alleles across all dogs within the DLA-88 locus. Similar trends were observed in DLA-88L, where although DLA-88*017:01 and DLA-88*029:01 exhibit the highest expression levels across all dogs, they are not the top two highly expressed alleles in Golden Retrievers. This phenomenon extends to other breeds such as Shih Tzus, Labrador Retrievers, Yorkshire Terriers, and others, indicating that alleles within DLA-88 and DLA-88L are breed-specific. Conversely, dominant alleles from DLA-12, DLA-64, and DLA-79 remain similar across all dogs and individual breeds, suggesting a lack of diversity within these loci.

**Figure 5.**
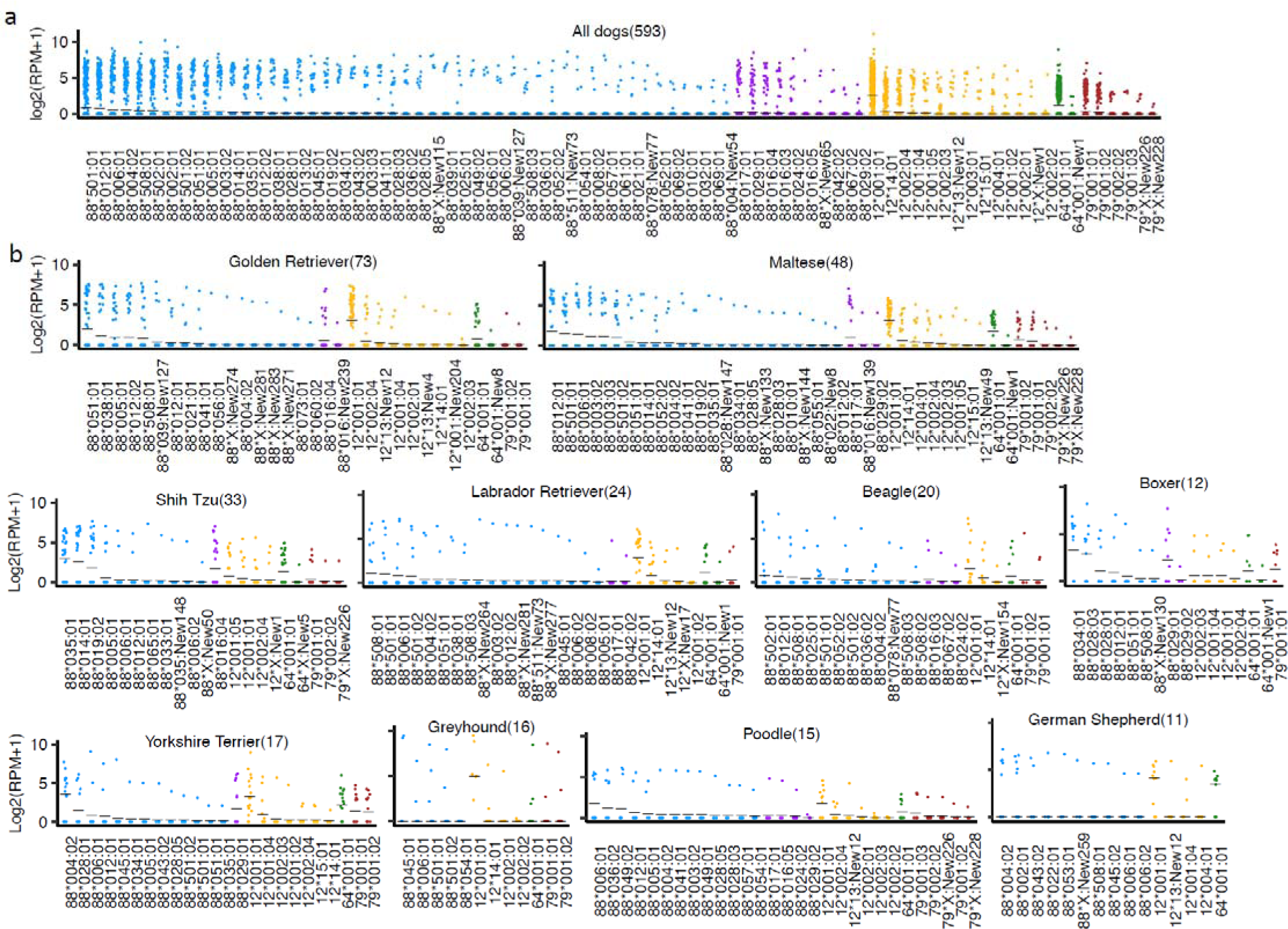
The dominant alleles across various breeds and the frequency of MHC-I alleles within the population. a. The RPM distribution of MHC-I alleles is arranged in descending order. Each point represents the log2(RPM+1) of the allele for each sample. The black dashes indicate the mean of log2(RPM+1) of each allele. The color coding represents different loci: DLA-88 in blue, DLA-88L in purple, DLA-12 in yellow, DLA-64 in green, and DLA-79 in brown. b. The prevalent alleles within 10 breeds (with a population size of ≥ 10 dogs) are determined based on log2(RPM+1) values without cutoff. The number in parentheses following each breed indicates the total number of dogs within that breed.

### MHC-I Diversity Comparison Between Human and Dogs

Expanding upon our earlier analyses, our investigation aims to explore the diversity of MHC-I genes between humans and dogs. As shown in Fig. 6a, distinct ethnicities present notable differences in the predominant expression of HLA-A, -B, and -C genes. Specifically, individuals of Caucasian descent typically express HLA-A*02:01 and HLA-A*01:01, while those of Asian descent commonly exhibit DLA-A*11:01 and DLA-A*24:02 (Fig. 6a). Each ethnicity showcases a unique tendency toward expressing specific alleles. Similarly, HLA-B and HLA-C also manifest varying dominant allele expression tendencies among different ethnic groups (Fig. 6b, c). Remarkably, while HLA-B does demonstrate a certain degree of tendency within each ethnicity, it lacks the pronounced tendency observed in HLA-A and HLA-C, indicating a higher degree of diversity in HLA-B alleles within and among ethnicities. Likewise, different dog breeds display distinct tendencies in DLA-88 expression. For instance, Golden Retrievers are predisposed to express DLA-88*051:01 and DLA-88*005:01, whereas Shih Tzus are more likely to express DLA-88*035:01 and DLA-88*014:01 (Fig. 6d). Overall, the diversity of DLA-88 expression within each breed is smaller compared to the diversity observed within each ethnicity across HLA-A, -B, and -C genes. However, the overall diversity observed in DLA-88 expression is similar to that observed in HLA-B expression (Fig. 6e). This suggests that DLA-88 in dogs exhibits a level of diversity comparable to that of HLA-B in humans.

**Figure 6.**
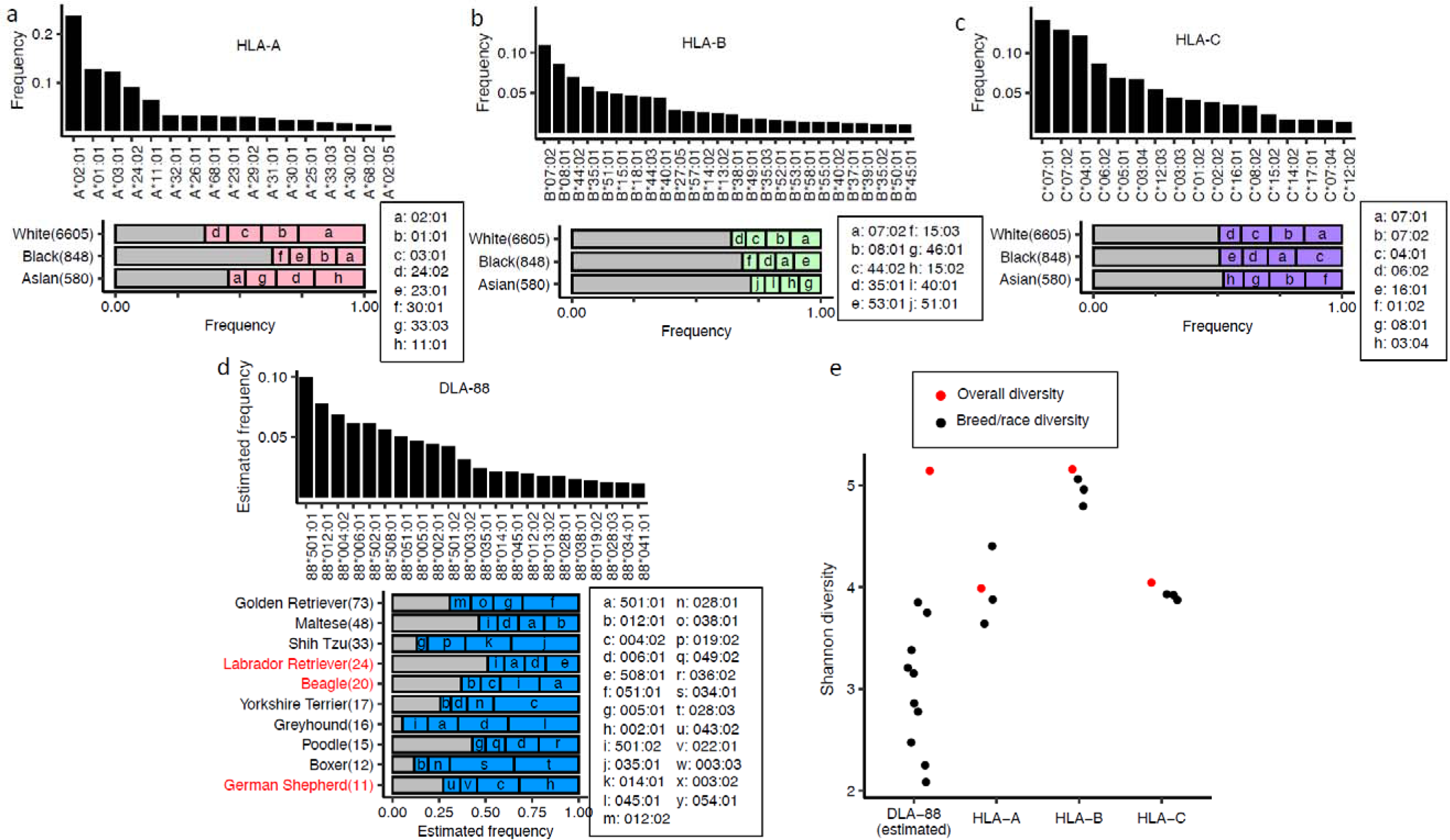
The comparison of MHC-I allele frequency and prevalent alleles in different populations between human and canine. a-c. The frequency of HLA-A (a), HLA-B (b), and HLA-C (c) alleles, along with the top four most common alleles, are presented for three human populations. The frequency distribution panels only display the alleles whose frequency >= 0.1%. Each letter corresponds to an allele. The sample size for each population is indicated in brackets next to the population name. d. The estimated allele frequency of DLA-88 alleles is presented in descending order, with the top four prevalent DLA-88 alleles identified within 10 breeds (each breed comprising ≥ 10 dogs). The sample size is indicated in parentheses. e. Diversity comparison across canine estimated allele frequency of DLA-88, HLA-A, HLA-A, and HLA-C. A dot represents a breed or a population. The red dots indicate the diversity of overall samples in canine or human.

### Dog and Human Share Hyper Variable Sites In MHC-I Potential Involved in Peptide Binding

We further investigated the amino acid composition of the exon2 and exon3 regions of human and dog MHC-I molecules, which are crucial regions implicated in peptide binding to MHC-I. Analysis of human MHC-I alleles, including HLA-A, -B, and -C, revealed varying degrees of amino acid diversity within the first 50 positions, with HLA-C exhibiting slightly higher diversity. However, overall, both human and dog MHC-I alleles did not exhibit high diversity within this initial segment. Notably, a strikingly high level of amino acid diversity was observed in regions spanning positions 75 to 100 and 155 to 165 across different species and within species (Fig. 7), suggesting shared hyper-variable regions between human and dog MHC-I molecules. Furthermore, examination of the amino acid composition in these regions revealed differences both across species (highlighted by red dots representing MHC-I human-dog comparisons in Fig. 7) and within species (indicated by green dots representing MHC-I variation across HLA alleles in Fig. 7). These positions with differing amino acid compositions significantly overlap with regions predicted to be in contact with peptides within the MHC-I binding cleft^22^, particularly in proximity to the B and F pockets, which are crucial for peptide binding.

**Figure 7.**
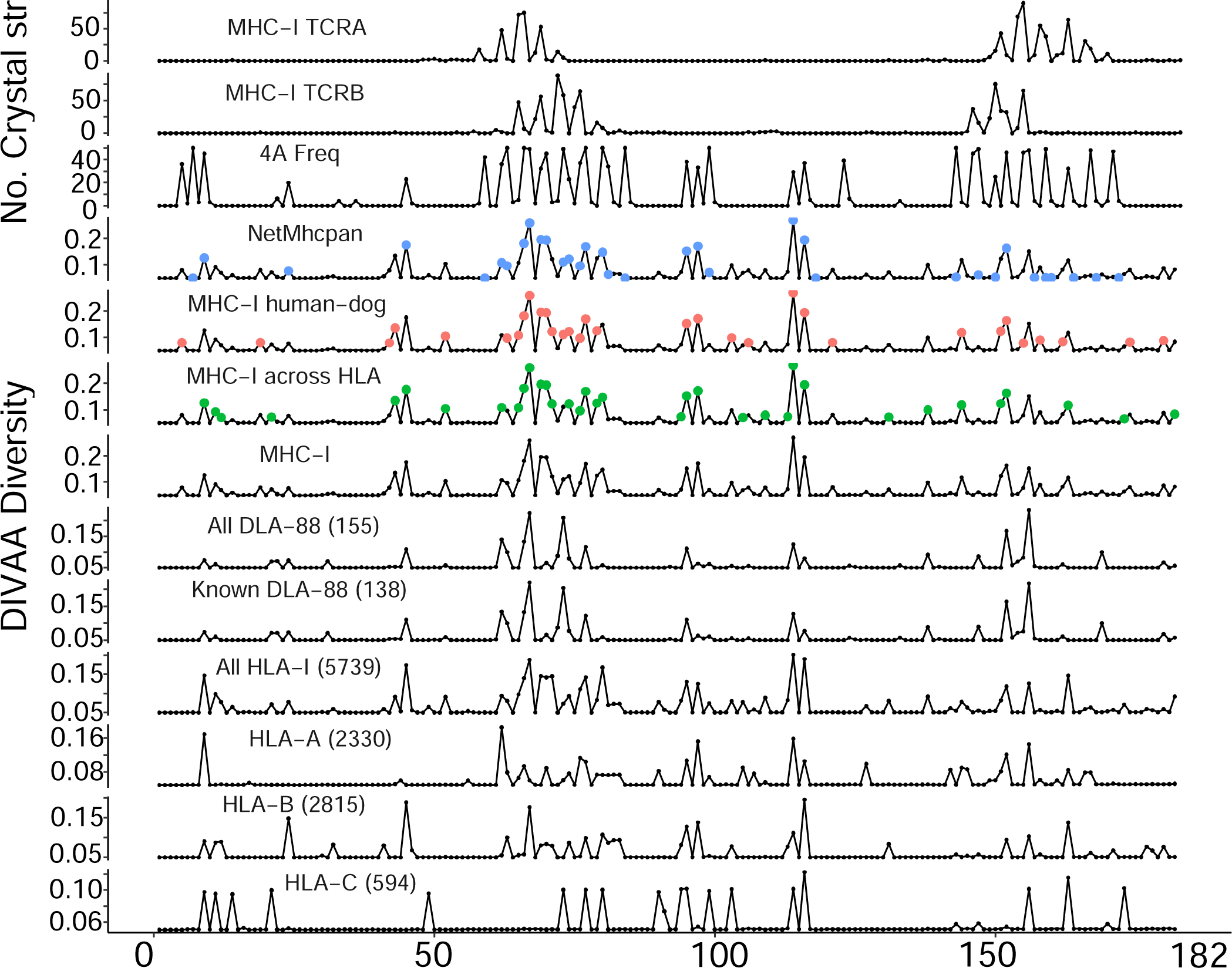
DIVAA diversity of MHC-I amino acids in human and dog indicates hyper variable sites across species. Each label above the line plot specifies the DLA or HLA alleles, with parentheses indicating the number of alleles included in the analysis. MHC-I includes both DLA-I and HLA-I alleles for the analysis (See also Methods). Green dots indicate the positions of MHC-I contacting binding peptides within 4 Å. Red dots represent positions with differing amino acids between humans and dogs. Blue dots indicate the positions with different amino acids among HLA-A/B/C. Purple dots indicate the positions HLA sequence estimated to be in contact with peptide in the binding cleft.

To complement our observations, we analyzed 457 MHC-I peptide binding complexes from the Protein Data Bank (PDB) and identified positions with a 4 Å distance between MHC-I molecules and binding peptides. Remarkably, these positions aligned closely with those around positions 75 to 100 and 155 to 165 in MHC-I exon2 and exon3 (Fig. 7, 4 Å Freq). Moreover, similar locations have been implicated in MHC-I and T cell receptor (TCR) binding, as evidenced by the binding site between TCR and MHC-I tending to localize within these regions (Fig. 7, MHC-I TCRA, TCRB).

In summary, our findings suggest that humans and dogs share similar hyper-variable sites in MHC-I molecules, and these sites may play crucial roles in both peptide recognition and TCR recognition.

## Discussion

### The identification of new alleles

The understanding of canine MHC-I diversity landscape was previously hindered by the lack of a high throughput approach to analyze the huge amount of public canine NGS data, and an effective way to identify new alleles. This situation has been improved, with the development of the KPR *de novo* assembler and genotyper ^11^ designed for canine MHC-I genotyping.

However, we also have to acknowledge that this kind of assembly-based approach has a higher requirement about the expression level of the MHC-I alleles, compared with some other best match searching-based software tools ^19,23–25^. And currently, the genotyping of alleles from some lowly expressed canine DLA-I genes is not as effective as genotyping DLA-88. So, this may influence the completeness of the landscape picture of genes like DLA-12, -64 and -79.

But on the other hand, the identification of new DLA-I allele candidates will in return, improve current situation in canine MHC-I genotyping, by improving the canine MHC-I read extraction in the algorithm, and bridging the gaps between human and canine MHC-I genotyping. Future experimental validation is required for the validation of all these candidate alleles.

Another goal of this study is to summarize all known DLA-I alleles. We noticed that there is some ambiguity in current DLA-I allele group definition and naming. Even the well-studied DLA-88 gene is facing this problem. For instance, DLA-88*019:01 and DLA-88*019:02, even though given the same allele group assignment, these two alleles have an amino acid difference in their first HVR.

### DLA-I allele frequency

Our genotyping identified a list of high frequency alleles across the 5 DLA-I genes. The result supports the previous conclusion that DLA-88 is the only classical MHC-I encoding gene in canine, while DLA-12, -64, and -79 show limited diversity and expression levels. DLA-88L exhibits a similar expression pattern to DLA-88 as it is not dominated by a single allele. But its sequence diversity is significantly lower than DLA-88. Considering its high sequence similarity to DLA-88, it is possible that some DLA-88L alleles were misclassified as DLA-88. And long-range sequencing is required, to understand the haplotype and linkage of this genomic region.

Most of the high frequency alleles match known alleles. But we are still able to identify some new alleles candidates with high frequencies, for instance, DLA-88*X:New115. One possible reason is that most of the known alleles are indeed high frequency alleles, which have a higher chance to be selected and studied in past research. Also, most of the previous canine MHC-I studies were conducted in some popular breeds, and these known alleles can also be the prevalent alleles in these breeds. We may have a different picture if more dogs from other geographic origins are incorporated into the analysis.

### MHC-I expression in different canine tissue types

The expression level of MHC-I molecules may be positively correlated with efficiency of immunotherapy. Therefore, understanding the transcription rate in different tissue types can help us predict clinical outcomes. However, almost all our samples belong to bulk RNA-seq, this measurement may be inaccurate if the cell type heterogeneity is high in the tissue. One possible example is the high expression observed in our B-cell and T-cell samples. Although MHC-I molecules are expressed in all nucleated cells, we could not find any publication supporting the up-regulation of MHC-I in these two cell types. This new finding is thrilling, but more in-depth studies are required in the future to root out the possibility of contamination caused by other cell types.

### The use of HVR in canine MHC-I classification

Finally, adding the newly found DLA-I alleles, we have refined HVRs of DLA-I, which currently are used to assign DLA-88 allele groups ^26^. Same HVRs can also be applied to define the allele groups in DLA-12 and -64 genes, as no significant difference was observed regarding this diversity between DLA-88 alone and all DLA-I alleles.

However, we note that HVRs also harbor sites that are not diversified, and some single highly diversified sites are located outside the HVRs. Hence, we propose that using “hyper-variable sites” (HVSs) may better capture the diversity of DLA-I alleles, leading to more accurate classification of DLA-I alleles and ultimately, prediction of peptide-binding specificity of DLA-I molecules.

## Material & Methods

### Data Collection

Canine RNA-seq data were retrieved from the Sequence Read Archive (SRA) database, encompassing diverse cancer types such as mammary tumor (PRJNA489087), oral melanoma (PRJNA749900), osteosarcoma (PRJNA525883), glioma (PRJNA579792), hemangiosarcoma (PRJNA562916), and others as detailed in Table S1. Additional information was sourced from relevant publications associated with these studies. The canine genome assembly (canFam3.1) and gene annotation (canFam3 1.99 GTF) were obtained from the Ensembl database. Allele frequency data for dogs were acquired from a pertinent study ^21^. For human MHC-I allele frequencies, data were obtained from a study which conducted HLA typing with OptiType ^27^ using 10,452 samples from TCGA ^28^. Further details are provided in Table human_HLA_allele_freq_NMDP_042623.xlsx.

### Canine RNA-seq data quality control (QC) and processing

RNA-seq read pairs were mapped to the canine reference genome canFam3 using HISAT2 (version 2.21)^29^ and concordantly and uniquely mapped pairs were reported by HISAT2. Subread (version 2.0.0)^30^ was used to calculate CDS-targeting rate with the number of reads with ≥1 bp overlapping a coding sequence (CDS) region of the canFam3 1.99 GTF. MultiQC (version 1.5)^31^ was used to examine GC content and duplicate level. The distributions of per sample read-pair total amount, mapping quality, and CDS targeting rate were examined to identify and exclude samples that fail to meet the cutoffs. A total of 65 RNA-seq samples failed the QC and were excluded from further analysis (Figures 1a-e). For each sample that passed QC measures, Cufflinks version 2.2.0^32^ were used to calculated FPKM (fragments per kilobase of exon per million mapped) value of each gene in each sample, which was then converted to TPM (transcript per million).

### Breed predictions and validations

We used 2,183 breed-specific variants identified as described^9^ in each dog sample to perform breed prediction and validation. Breed validation was achieved via standard hierarchical clustering with VAF values of breed-specific variants of each dog. If dogs contain both normal and tumor samples, the mean of the VAF values was used. The figure is illustrated in Fig. 1f.

### DLA-I genotyping by KPR

MHC-I genotyping was performed on both internal and public canine bulk RNA-seq samples using KPR^11^. For utilizing this software, we set the Kmer length (K) to half the sequence length of each sample and conducted 1,000 rounds of assembly runs (N). Default read pair depth (PD) values (PD=15 for DLA-88 and PD=38 for DLA-12) were employed for contig validation.

### New allele discovery and validation

The KPR output from each sample was utilized to evaluate DLA-I alleles. Contigs obtained from KPR with a “Long identical head/tail” label in the confidence column were excluded from the analysis to reduce false positives, except for those also reported in ^21^. Each contig was categorized based on the following criteria: contigs with a relative expression level greater than 25% (Rel >25%), contigs within the top four highest relative expression levels within a sample (Top 4), contigs identified in more than three dogs (Freq ≥ 3), contigs found in the known DLA-I database (Known), and contigs detectable with Seq2HLA^19,20^ in the same sample (Seq2HLA). Only alleles meeting at least one of these conditions were classified as validated alleles. For DLA-79, each allele identified in KPR was manually validated. A phylogenetic tree was constructed using the Neighbor-Joining method and assessed through 10,000 bootstrap replicates. This was achieved after aligning the protein sequences of exon2 and exon3 of DLA-I using MEGA 11 software, which can be accessed at https://www.megasoftware.net/ (accessed on July 3, 2023).

### DLA-I tissue-specific differential expression

DLA-I reads mapped to DLA-I genes were used to analyze the differential tissue expression. The DLA-I reads normalized against each sample’s total reads to plot in Figure 4. We used the reads ratio for each DLA gene to estimate DLA reads in DLA-88, DLA-88 50X, DLA-64, and DLA-79. Sequencing evenness was assessed with Kolmogorov-Smirnov test between the actual read coverage distribution in exon2 and exon3 regions and the theoretical uniform distribution.

### Correlation between genotyping rate and total DLA exon2-3 sequence read pairs or sequence error rates

A median of DLA exon2-3 read pairs or median of sequence error rates of all passed QC samples was used for each BioProject. The genotyping rate is the ratio of samples with at least one DLA genotyping result to the passed QC samples of each BioProject.

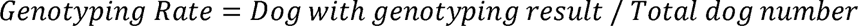

### Genotyping Rate = Dog with genotyping result / Total dog number

Sequence error rates are defined as total mismatched base pairs divided by total base pairs in each sample. The Pearson correlation and Spearman correlation were performed to identify the correlation coefficient. To combine total DLA exon2-3 read pairs and sequence error rates, we performed multiple linear regression to get the weight for DLA exon2-3 reads pairs (Wd) and sequence error rates (Ws) and intercept (I). To combine DLA exon2-3 read pairs and sequence error rates as a new combine variable (C), we use

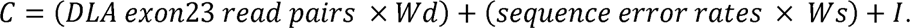

And the exon2-3 sequence uniformity test was conducted by the SciPy package using the coverage at each base position in exon2-3 region.

### Sanger Sequencing

PCR cloning-based Sanger sequencing was performed for internal canine samples. The sequencing results were genotyped with a standalone pipeline:

● Extraction of Sanger sequences with complete exon2-3 coverage.
● Sequence translation.

Comparison with database alleles to achieve a 5-digit typing, or identification of the closest database if no perfect match was found.

### Breeds clustering with non-negative matrix factorization

“The NMF R package^33^ was applied to process canine samples, specifically those containing DLA-88 and DLA-88L expression levels identified from KPR. The expression levels were normalized to reads per million (rpm) units. To determine the optimal rank for the analysis, 100 iterations were performed. To minimize the impact of noise signals, we include breeds with a sample size of 10 or more dogs for the NMF clustering. Breed enrichment analysis was performed using the basis gene lists identified by the NMF.

### MHC-I hypervariable sites identification

5739 DLA-I sequences were downloaded from http://hla.alleles.org/ (02/17/2022). 184 DLA-I sequences were identified in this study. To make DLA and HLA the same length, we inserted one extra empty amino acid sequence in 155 in DLA and HLA alleles except in DLA 50X alleles. DIVAA diversity, as described in^34^ was applied to analyze the amino acid diversity for each position. To account for the different amounts of HLA alleles in different genes, we repeat 100 times to perform random sampling for HLA genes comparison (All HLA-I) and repeated 10000 times for DLA-HLA(MHC-I) comparison and use the mean values to plot the result. To identify the differences in the amino acid of MHC-I position among HLA (MHC-I across HLA) and between DLA and HLA (MHC-I human-dog), we created consensus sequences (the most frequent sequence) for HLA-A/B/C and DLA and highlighted the position as illustrated in Figure 7.

### Estimated Allele Frequency

We classified each dog sample as homozygous or heterozygous for a specific locus and counted the number of alleles carried by each dog sample. A dog was considered homozygous if it carries one allele for the locus and heterozygous if it carries two or more alleles. The total number of alleles in the population was calculated as follows:

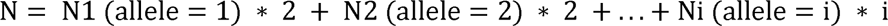

Where N1 represents the number of homozygous dog samples, and Ni represents the number of heterozygous dog samples carrying i alleles. The number of the specific allele of interest was calculated as:

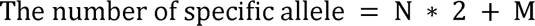

Where N is the number of homozygotes carrying this allele, and M is the number of heterozygotes carrying this allele. The estimated allele frequency was then determined by dividing the number of the specific allele of interest by the total number of alleles in the population.

## Declarations

None

### Ethics approval and consent to participate

Not applicable.

### Consent for publication

Not applicable.

### Author contributions

All authors conducted the analysis, wrote, and approved the manuscript.

## Acknowledgments

We thank Dr. Jingxuan Chen for providing help on the breed validation analysis; the Georgia Advance Computing Resource Center, University of Georgia (GACRC) for supporting this work. This work is funded by NCI R01 CA252713, R01 CA182093 and AKC Canine Health Foundation.

